# MultiSAAl: Sequence-informed Antibody-Antigen Interactions Prediction using Multi-scale Deep Learning

**DOI:** 10.1101/2025.05.29.656915

**Authors:** Zexin Lv, Dongliang Hou, Minghua Hou, Suhui Wang, Jianan Zhuang, Guijun Zhang

## Abstract

**Motivation:** Antibodies possess high specificity toward target antigens, making them critical for therapeutic applications. However, experimental screening of antibody-antigen interactions is labor-intensive and costly. Existing structure-based computational methods often face significant challenges in modeling the dynamic conformational flexibility that governs antibody-antigen interactions, which limits their ability to accurately identify binding. Therefore, there is an urgent need for accurate and efficient sequence-based prediction method to accelerate antibody discovery and reduce experimental burden.

**Results:** We propose MultiSAAI, a sequence-informed framework that models antibody-antigen interactions by explicitly accounting for the distinct roles of antibody heavy and light chains in antigen binding. MultiSAAI integrates language model embeddings, physicochemical properties, geometric constraints, and residue substitutability to characterize antibody-antigen interactions across multiple scales, employing a multi-scale network architecture that simultaneously evaluates global residue-pair compatibility and local amino acid fitness at the binding interface. Furthermore, the incorporation of site-specific information and biologically grounded binding principles allows the model to more closely reflect the actual mechanisms of interactions. Benchmark results demonstrate that MultiSAAI achieves AUROC scores of 0.757 on the generic antibody-antigen interactions dataset and 0.946 on the SARS-CoV-2 dataset, outperforming existing methods such as A2binder and AbAgIntPre. Finally, large-scale preliminary antibody screening further validates the potential of MultiSAAI for high-throughput therapeutic antibody discovery.

## 1 Introduction

Antibody-antigen interactions (AAIs) are critical to immune defense, enabling antibodies produced by B cells to selectively recognize and neutralize pathogens such as bacteria and viruses. Beyond their natural immune roles, AAIs serve as the foundation for diverse biomedical applications, ranging from diagnostic assays and therapeutic interventions (Scott, et al., 2012; Andrews, et al., 2022; Watanabe, et al., 2023) to structure-guided vaccine design (Steichen, et al., 2019). The characterization of AAIs represents a fundamental challenge in immunology and therapeutic development. Traditional experimental techniques, including phage display (Ledsgaard, et al., 2018) and enzyme-linked immunosorbent assays (ELISA) (Grange, et al., 2014), while providing valuable binding information, suffer from critical limitations in throughput, scalability, and cost-effectiveness. These persistent challenges in experimental AAIs characterization have driven an urgent demand for alternative approaches in therapeutic antibody development. To address these challenges, computational approaches have revolutionized AAIs research by overcoming key experimental limitations through enhanced efficiency, scalability, and cost-effectiveness.

Currently, computational methods for AAIs prediction are primarily categorized into two groups: structure-based and sequence-based approaches (Kim, et al., 2023). Structure-based methods can be categorized into two main approaches: one that employs energy-based scoring functions to assess antibody-antigen docking (Pierce, et al., 2011; Sulea, et al., 2016) and another that uses deep learning to predict the accuracy of antibody-antigen interactions or docking poses(Schneider, et al., 2022; Myung, et al., 2022). However, both the prediction of antibody-antigen binding and the scoring of docking depend on high-precision structures, and the extensive experimental effort needed to obtain these structures significantly limits the practicality of these methods. Moreover, structure-based predictions face three major challenges. First, antibody binding sites are primarily composed of complementarity-determining regions (CDRs) (Gabrielli, et al., 2009), which exhibit a diverse range of amino acid arrangements and three-dimensional conformations that conventional protein interactions models cannot fully capture (Adolf-Bryfogle, et al., 2015). Second, antigenic epitopes show significant diversity from continuous linear sequences to discontinuous conformational motifs (Vita, et al., 2019), each of which affects antibody binding in a unique way, adding difficulty to identifying antigenic epitopes that interact with antibodies in the interactions. Third, the dynamic flexibility of the CDRs facilitates essential conformational adaptations during antigen recognition (MacCallum, et al., 1996); while traditional rigid docking simulations struggle to represent this process adequately, complicating the evaluation of interactions based solely on the docking pose.

The widespread adoption of sequence based computational methods for antibody–antigen interactions prediction primarily stems from the greater availability and experimental tractability of sequence data compared to structural information. One representative methodology involves employing convolutional neural network (CNN) to analyze a large collection of diverse antibody sequences, thereby enabling the screening of antibodies that specifically recognize HER2 (Mason, et al., 2021). AbAgIntPre (Huang, et al., 2022) utilizes a Siamese-like CNN that uses k-spaced amino acid pair (CKSAAP) encoding to capture sequence features of both antigens and antibodies, thereby enabling accurate binding prediction. Similarly, DeepAAI (Zhang, et al., 2022)combines adaptive relation graph learning to capture global interactions patterns with CNNs that extract local sequence features for predicting antibody-antigen neutralization effects. The development of protein and antibody language models, including ESM-2 (Lin, et al., 2023), ProtTrans (Elnaggar, et al., 2021), AbLang (Olsen, et al., 2022), and AntiBERTy (Ruffolo, et al., 2021), have demonstrated remarkable capabilities in biomolecular interactions prediction. These models effectively leverage large-scale sequence data to infer structural information, thereby enabling sequence-based approaches that relieve the limitations of structural data. For example, A2binder (He, et al., 2024) collected over one billion antibody light and heavy chain sequences from the Observed Antibody Space (OAS) database (Olsen, et al., 2022) to train Roformer models for both chains, thereby further enhancing binding affinity prediction. Despite these advances, existing sequence-driven frameworks tend to focus on analyzing global sequence information without fully leveraging the key interactions regions, such as the antibody binding sites (paratopes) and antigen binding sites (epitopes). Recently, DeepNano (Deng, et al., 2024) leverages antigen-binding sites in antibody-antigen interactions to improve prediction accuracy, while AntiBinder (Zhang, et al., 2025) emphasizes the CDRs to aid in interactions prediction. Both methods are forward-looking and offer valuable insights into the potential mechanisms governing antibody-antigen interactions.

To bypass the requirement for structural data in antibody-antigen interactions prediction while improving prediction performance, we propose a multi-scale deep learning framework, MultiSAAI, that establishes a sequence-based paradigm for accurate interactions prediction. MultiSAAI employs a bilinear attention network (BAN) (Kim, et al., 2018) to model pairwise residue interactions, coupled with a multi-fusion convolutional neural network (MF-CNN) (He, et al., 2024) that processes interactions across different spatial scales. The architecture integrates both antigenic epitopes and CDR-H3 information, which have been structurally validated as critical binding determinants (Marks and Deane, 2020), to enhance prediction accuracy. Experimental results show that MultiSAAI outperforms existing methods across diverse antigen types and offers interpretable insights into predicted binding outcomes. Furthermore, MultiSAAI has been validated in antibody screening experiments, demonstrating its capability to accurately identify antibodies that bind to specific antigen targets.

## 2 Method

### 2.1 Dataset

In this study, we developed two deep learning models, MultiSAAI and MultiSAAI_SARS2, which share identical network architectures and feature engineering pipelines. The key difference between these two models lies in their training datasets. MultiSAAI was trained on our newly constructed generic dataset of antibody-antigen interactions derived from SAbDab (Dunbar, et al., 2014), while MultiSAAI_SARS2 was trained and evaluated using the same dataset as A2binder (He, et al., 2024) to enable direct performance comparisons for SARS-CoV-2 variant binding predictions. This dual-dataset was strategically designed to: (1) validate model generalizability through our comprehensive, newly curated dataset covering diverse antibody-antigen interactions, and (2) ensure fair benchmarking against existing methods (A2binder) on established SARS-CoV-2 data, thereby providing both broad applicability assessment and focused performance validation on a clinically critical target.

The MultiSAAI dataset was constructed through a multi-stage protocol to ensure biological validity and minimize data redundancy. From the SAbDab (July 2024 release), we extracted 4440 complete protein antibody-antigen complexes containing paired heavy chain variable, light chain variable, and their corresponding antigen sequences. Redundant antibody sequences were removed using CD-HIT (Fu, et al., 2012) with a 98% sequence identity threshold, yielding 2609 non-redundant interactions pairs. For positive sample generation, antigen sequences were clustered into 747 groups with 90% identity using MMseqs2 (Steinegger and Söding, 2017), followed by two data augmentation strategies: intra-group antibody random pairing and CDR-H3-restricted antigen exchange, which collectively generated 3790 positive samples (including 1181 newly added positive samples). Negative samples were constructed through stringent inter-group pairing to maintain a balanced 1:1 ratio between positive and negative samples. To preserve evolutionary diversity and prevent data leakage, we performed phylogenetic clustering on all 7580 samples using ClustalW (Thompson, et al., 2003), identifying nine evolutionarily distinct clusters that were each split at a 4:1 ratio into training and test sets. Additional details are provided in **Supplementary Note 1.**

### 2.2 The MultiSAAI Pipeline

The pipeline of MultiSAAI is illustrated in **Fig.1**. The model takes as input the amino acid sequences of both the heavy and light chains of an antibody along with the corresponding antigen sequence, and generates as output a predicted interaction score. MultiSAAI is a specifically designed deep learning framework that effectively integrates multi-scale features with a Multi-scale Interactive Learner (MIL) for accurate antibody-antigen binding prediction.

**Figure 1.**
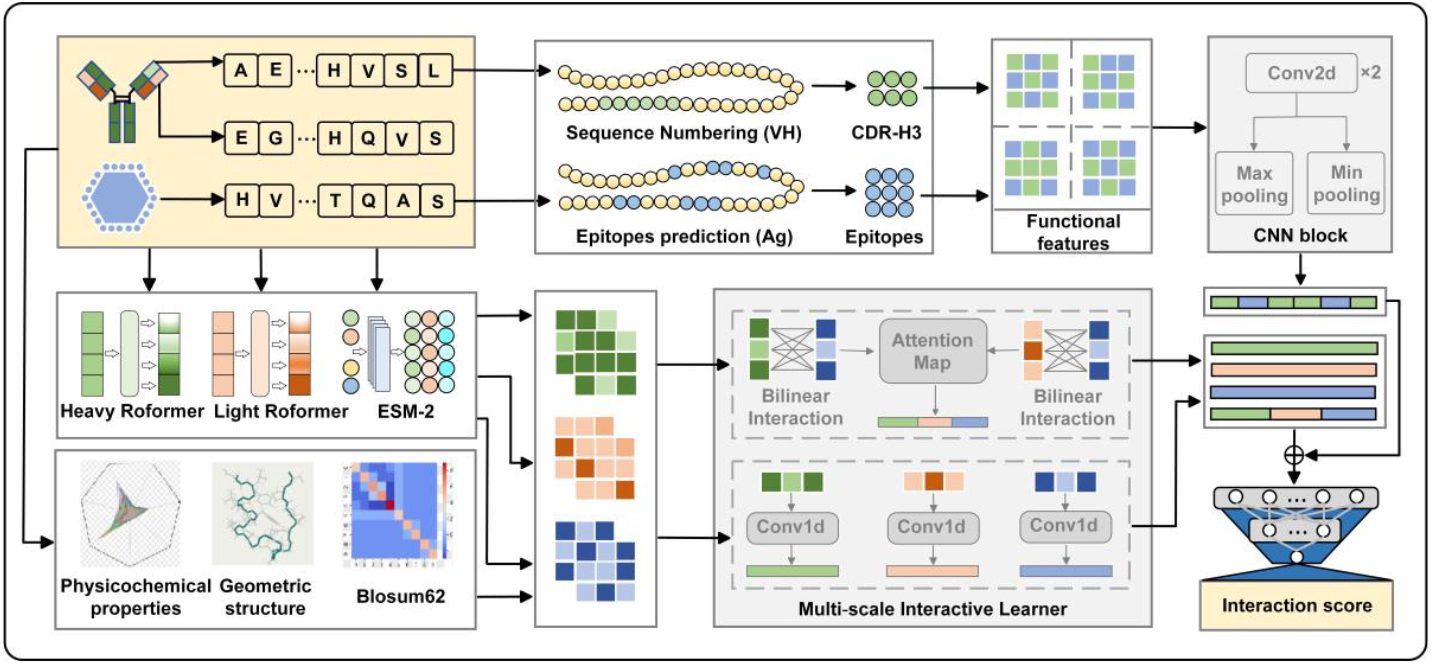
MultiSAAI takes antibody light chain, heavy chain and antigen sequences as input. The Sequence semantic features, Physicochemical features, Geometric features, and Evolutionary features of the three sequences are learned through the Multi-scale Interactive Learner. At the same time, the proprietary CNN block processes the Functional features and finally outputs the interaction score.

The multi-scale feature representation system comprises five characteristic dimensions: (i) sequence embeddings from protein language models, (ii) residue-specific physicochemical properties, (iii) secondary structure geometric constraints, (iv) evolutionary conservation patterns, and (v) functional interactions signatures. Notably, the functional interactions signatures (i.e., functional features) specifically highlight the characterization of critical interactions sites. By focusing on the molecular recognition mechanisms between antibody CDR-H3 hypervariable loops and antigenic epitopes, the engineered features achieve high-fidelity modeling of antibody-antigen interactions interfaces. This integrated multi-scale feature strategy not only comprehensively captures all critical determinants of antibody-antigen binding interactions but also significantly enhances predictive accuracy by optimizing representation at key functional sites.

The Multi-scale Interactive Learner (MIL) network architecture, which integrates a Bilinear Attention Network (BAN) and a Multi-Fusion Convolutional Neural Network (MF-CNN), is specifically designed to generating a global interaction matrix that captures inter-molecular binding patterns. The MF-CNN module, which processes antibody and antigen sequences through hierarchical convolutional layers with varying receptive fields, extracts local features that are critical for binding specificity. Notably, this architecture effectively eliminates feature contamination artifacts induced by direct sequence concatenation between heavy chain and light chain while preserving strict separation of molecular representations, thereby enabling comprehensive cross-modality interaction analysis. Additionally, the functional features undergo dedicated convolutional module to enhance the characterization of local-scale interactions between CDR-H3 and epitopes.

### 2.3 Feature Extraction

#### Functional features

Given the critical role of the heavy chain CDR-H3 in epitope binding and the fundamental contribution of noncovalent forces to antibody-antigen interactions (Liu, et al., 2024), we developed functional features to quantitatively characterize this region. These interaction features are structured as a contact matrix that comprehensively captures noncovalent interactions between CDR-H3 and epitope residues, including hydrogen bonds, ionic interactions, van der Waals contacts, and hydrophobic effects. CDR-H3 residues were identified and numbered using ANARCI (Dunbar and Deane, 2016) following the IMGT unique numbering scheme (Lefranc, et al., 2009). Antigenic epitopes were predicted using our in-house developed epitope prediction tool (currently in development, with public release planned upon completion). Detailed information on the functional feature is also provided in **Supplementary Note 2**.

#### Sequence semantic features

Sequence semantic features were extracted from protein sequence language model embedding, leveraging the pre-trained models’ ability to capture high-order semantic relationships within and between antibody-antigen sequences. Specifically, Roformer processes antibody sequences while ESM-2 (Lin, et al., 2023) handles antigen sequences (He, et al., 2024), with this specialized division leveraging each model’s unique strengths. Roformer is selected for antibodies due to its ability to efficiently capture long-range dependencies, which are critical given the complex folding and interaction patterns of antibodies. ESM-2 is used for antigens to leverage its strong generalization ability and pretraining on diverse protein sequences, thus enabling accurate modeling of antigen sequence properties.

#### Physicochemical features

Physicochemical features include normalized sequence end distance and descriptors reflecting amino acid-specific chemical properties (Meiler, et al., 2001), such as hydrophobicity, charge, and polarity. These features describe the local positional environment and chemical characteristics of each residue, providing important complementary information beyond sequence semantics for the prediction of antibody-antigen interactions.

#### Geometric features

Geometric features characterize the influence of secondary structures on antibody– antigen interactions. Residues located within specific structural motifs, such as helices, sheets, and coils, often display distinct patterns of solvent accessibility and spatial organization, which in turn affect their likelihood of participating in binding. Capturing secondary structure-related geometry enhances the model’s ability to infer interaction-relevant spatial proximities.

#### Evolutionary features

While sequence semantic features from language models capture sequence-dependent patterns in antibody-antigen pairs, evolutionary features provide sequence-independent constraints derived from the BLOSUM-62 substitution matrix (Eddy, 2004). Blosum-62 substitution matrix quantifies the likelihood of amino acid substitutions based on evolutionary conservation. This information highlights residues that are functionally or structurally important, providing additional cues for identifying critical binding sites between antibodies and antigens. Further details on features are summarized in **Supplementary Table S1**.

### 2.4 Network Model Architecture

#### Multi-scale Interactive Learner (MIL)

The MIL is architected using Bilinear Attention Network (BAN) in combination with multi-fusion convolutional neural network (MF-CNN), emphasizing the integration of both global structure context and local amino acid interactions. BAN, originally developed for multimodal tasks, is effective at capturing correlations between different feature types. In this study, we adapt the BAN architecture for antibody-antigen interactions (AAIs) prediction by computing pairwise attention scores between all residue pairs across antibody and antigen sequences, thereby explicitly modeling their binding. Similarly, MF-CNN effectively utilizes multiple convolutional kernels to integrate local embedding features, allowing it to identify detailed interaction patterns between neighboring residues.

BAN consists of two layers. The first layer uses a bilinear interaction mapping to compute attention weights for each pair of residues. The second layer applies a pooling operation to aggregate these weights into a comprehensive representation of the full sequence. All features except the functional features generate respective embeddings for the antibody heavy chain, light chain and antigen sequence as the input of the Bilinear Attention Network.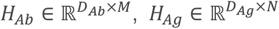 Where *H*_*Ab*_ and *H*_*Ag*_ are antibody embeddings and antigen embeddings respectively, *D*_*Ag*_ and *D*_*Ab*_ are their respective dimensions,820 and 692, *M* and *N* correspond to the sequence length. When the antibody embedding and antigen embedding are input into the Bilinear Attention Network, the MF-CNN captures local correlations among amino acids within *H*_*Ab*_ and *H*_*Ag*_. The module has three layers, including convolution, pooling and ReLU activation, to process the sequence embeddings. It then uses three linear layers with residual connections to combine these features and generate the final output representation.

#### CNN block

The functional features between CDR-H3 and the epitopes capture local interactions among a limited number of amino acids. Therefore, it is reasonable to process these local features using a CNN block. The input features have dimensions *L*_*e*_×*L*_*h*_×5. Here, *L*_*e*_represents the sequence length of the antigenic epitopes (in amino acids), *L*_*h*_represents the CDR-H3 loop amino acid length. After passing through two convolutional layers and applying both max and min pooling, to capture the salient information in the combined features.

#### Classifier

The classification module is a multi-layer perceptron (MLP) comprising three linear layers. Vectors generated by the Multi-scale Interactive Learner and the CNN block, are concatenated and fed into the MLP to output the interaction score. The detailed description of the MultiSAAI network model architecture is provided in **Supplementary Note 3**.

### 2.5 setting

During training, we fine-tuned the pre-trained language models for both antibodies and antigens, employing distinct learning rates to optimize the language models and network parameters, where the language models use a smaller learning rate (2e-5) to stabilize fine-tuning. We adopted Binary Cross Entropy (Ruby and Yendapalli, 2020) as the loss function, and all models were trained on NVIDIA A100 Tensor Core GPUs. Detailed parameter settings are provided in **Supplementary Note 4**.

## 3 Result

### 3.1 Performance of MultiSAAI

We compared MultiSAAI with two methods: A2binder (He, et al., 2024) and AbAgInPre (Huang, et al., 2022). A2binder was originally designed for SARS-CoV-2 interaction prediction and has excellent performance, while A2binder_SAbDab was retrained on the generic antibody-antigen dataset using its default parameters. AbAgInPre serves as a comparable baseline in our study, offering both a general-purpose model and a specific model that included SARS-CoV-2, which is consistent with the study tasks of MultiSAAI and MultiSAAI_SARS2. For an unbiased performance assessment with AbAgInPre, we conducted all comparative evaluations through the publicly available web server with consistent settings.

Compared to these two methods, our approach achieves superior overall performance and demonstrates enhanced accuracy in modeling the distribution of predicted interaction score. On the generic dataset, MultiSAAI consistently ranks highest across all evaluation metrics, reflecting its ability to robustly distinguish positive and negative samples **(Fig.2a)**. MultiSAAI exhibits a pronounced unimodal distribution of prediction scores in both positive and negative sample tests **(Fig. 2b)**, despite a small fraction of misclassified samples. This phenomenon primarily stems from the high sequence conservation in antibody variable regions, which biases the model toward overfitting certain recurrent structural patterns while potentially overlooking functionally important subtle variations. In comparison, A2binder_SAbDab on the same data performs poorly on a subset of pairs and identifies only a few true interactions. AbAgIntPre also fails to separate the two classes, clustering most predictions between 0.4 and 0.8. These findings highlight the challenge faced by existing models in capturing fine-grained interaction features, while underscoring the effectiveness of MultiSAAI in learning antibody–antigen binding representations.

**Figure 2.**
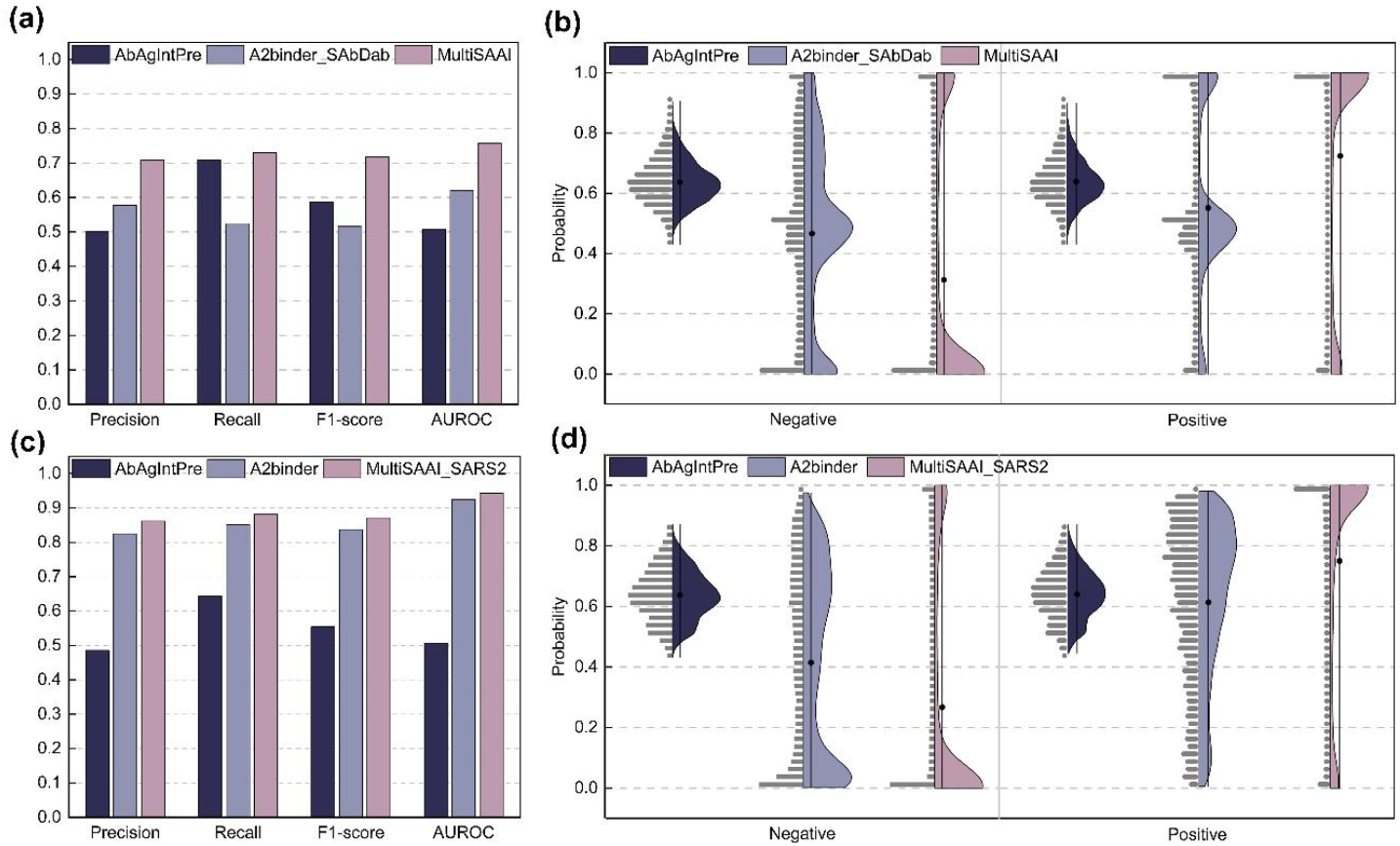
Figure (a) and Figure (b) represent the overall metrics and positive and negative sample tests of MultiSAAI and other methods. Figure (c) and Figure (d) represent the overall metrics and positive and negative sample tests of MultiSAAI_SARS2 and other methods.

When tested on the SARS-CoV-2 dataset, all models benefit from the increased data richness, leading to generally improved performance **(Fig. 2c)**. Although A2binder narrows the performance gap with MultiSAAI_SARS2 in overall metrics, the positive and negative sample tests reveal that MultiSAAI_SARS2 achieves more accurate distinction between positive and negative interactions **(Fig.2d)**. Meanwhile, AbAgIntPre again struggles to clearly resolve the two classes, with predictions concentrated in a narrow, uncertain range. These findings highlight that MultiSAAI and MultiSAAI_SARS2 effectively captures the binding rules governing antibody-antigen interactions. Detailed results for various metrics are provided in **Supplementary Tables S2-S5**.

### 3.2 Ablation study

We conducted ablation analysis on both MultiSAAI and MultiSAAI_SARS2 (**Fig.3a and Fig.3b**). We assessed the performance changes resulting from the stepwise addition of key components to the base model (v1). The v1 configuration consists of MF-CNN and ESM-2 applied to both antibody and antigen sequences. Based on this, v2 introduces physicochemical, geometric, and evolutionary features; v3 adds the Bilinear Attention Network (BAN); v4 replaces ESM-2 on the antibody side with the antibody Roformer; and v5 incorporates functional features.

**Figure 3.**
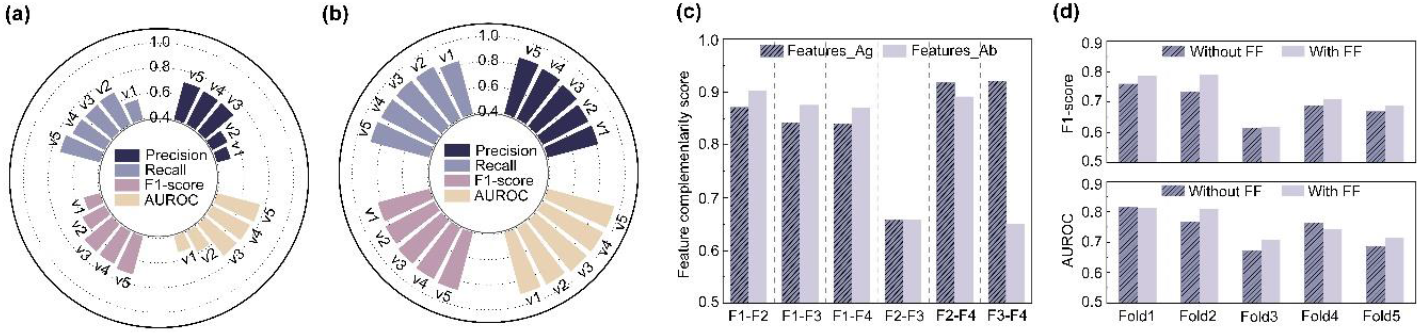
(a) Ablation experiments on MultiSAAI. (b) Ablation experiments on MultiSAAI_SARS2. (c) The complementarity analysis of each pair of features of antigen and antibody was performed, where F1, F2, F3, and F4 represent their respective sequence semantic, evolutionary, physicochemical, and geometric features. (d) Results of five-fold cross validation with and without functional features.

In the MultiSAAI ablation study, each added component contributes positively to performance. The most significant performance enhancement occurs between versions v2 and v3, where the sole architectural difference is the incorporation of the Bilinear Attention Network (BAN) module. This substantial gap highlights the importance of BAN for capturing fine-grained residue-level interactions between antibodies and antigens. Additionally, the inclusion of physicochemical, geometric, and evolutionary features effectively complements sequence information, enhancing interaction prediction. Although replacing ESM-2 with the antibody Roformer yields modest gains, the limited dataset size may constrain the full potential of language model refinements. Notably, the addition of functional features in v5 consistently improves all evaluation metrics, underscoring the critical role of noncovalent interactions between CDR-H3 and antigenic epitopes in interaction prediction.

For MultiSAAI_SARS2, each component similarly contributes incremental improvements in F1 and AUC scores. However, these gains are relatively small, likely due to the dataset’s richness. With 32 antigens and over 27,000 antibody sequences, the SARS-CoV-2 dataset provides ample diversity, allowing the model to learn robust interaction patterns even without auxiliary features. In contrast, the smaller scale of the generic dataset limits model performance, explaining the performance gap between MultiSAAI and MultiSAAI_SARS2 (**Supplementary Note 5**). Detailed ablation results are provided in **Supplementary Tables S6-S7**.

### 3.3 Complementary Feature Analysis

We evaluated the complementarity among the selected features for the antibody and antigen feature sets separately to ensure they capture non-redundant information. A higher complementarity score indicates lower overlap between features, supporting the diversity of the feature representation (**Fig.3c)**. Except for the sequence semantic features, which differ due to the use of different language models. Here, F1, F2, F3, and F4 represent sequence semantic, evolutionary, physicochemical, and geometric features, respectively. Overall, we observed that the complementarity between ESM-2 and other features is slightly lower than that between the antibody Roformer and other features. This finding supports our earlier experimental results, suggesting that the Roformer is more suitable for encoding antibody sequences than ESM-2. Detailed results on feature complementarity are given in **Supplementary Table S8**.

Interestingly, physicochemical and geometric features exhibit higher complementarity on the antigen side compared to the antibody side. This may reflect a greater structural flexibility of antigens, where secondary structure formation is less dependent on intrinsic amino acid properties. In contrast, antibody structure may rely more strongly on physicochemical characteristics such as hydrophobicity, charge, and polarity. These observations suggest that, while language models capture rich implicit sequence-level information, traditional features based on evolutionary, physicochemical, and geometric features remain valuable for enhancing the overall representation.

The functional features designed around the interactions between CDR-H3 and antigenic epitopes represent an innovation of our method. These features capture localized, noncovalent interactions patterns that are not easily represented by conventional residue-level descriptors. Although their two-dimensional nature prevents direct complementarity analysis like other features, we assessed their effectiveness through five-fold cross-validation (**Fig.3d)**. Incorporating these features generally improves model performance, with notable gains in both F1-score and AUROC, particularly in Fold 2. The modest decline in AUROC observed in Fold 1 and Fold 4 may be due to variability in epitopes prediction accuracy. These results suggest that when epitopes localization is reliable, functional features derived from the spatial relationship between CDR-H3 and antigenic epitopes can significantly enhance the model’s ability to capture meaningful interaction patterns. This highlights the biological relevance and practical value of incorporating domain-specific binding priors into deep learning frameworks, especially in scenarios with limited data. The detailed results of the contribution of functional features to MultiSAAI are in **Supplementary Table S9**.

### 3.4 Case Study: MultiAAI Screens the Potential Antibodies Targeting HER2

Screening candidate antibodies with strong antigen-binding capacity is a critical step in antibody drug discovery. HER2 (human epidermal growth factor receptor 2) is a well-established therapeutic target, particularly in breast and gastric cancers (Rubin and Yarden, 2001), due to its role in tumor progression through mutation or overexpression. In this study, we applied MultiSAAI to predict interaction score between HER2 and all antibody sequences targeting protein antigens from the SAbDab. The goal was to assess whether, under a pathogen invasion scenario, binding-competent antibodies could be rapidly prioritized using only the antigen sequence, thereby reducing experimental screening workload.

The distribution of predicted scores indicates that most antibodies are unlikely to bind HER2 (**Fig.4)**, which aligns with known biological expectations. Notably, trastuzumab and pertuzumab (Scheuer, et al., 2009), two monoclonal antibodies known to target HER2, received extremely high binding scores (0.995 and 0.998, respectively), even though their sequences were not included in the model training set. This finding supports the generalizability of MultiSAAI and its potential to identify therapeutically relevant antibodies with high confidence. While a small number of false positives were observed, the overall results demonstrate the model’s utility in large-scale virtual screening and suggest its promise in accelerating early-stage antibody discovery for specific antigens.

**Figure 4.**
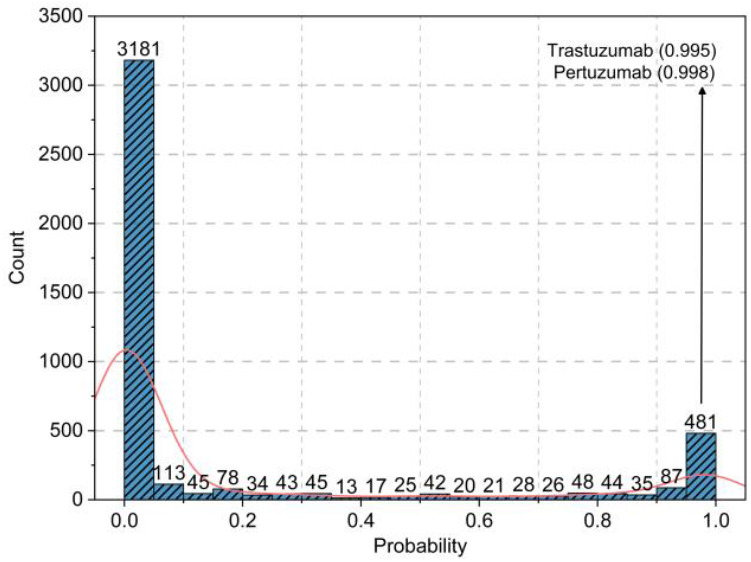
We applied MultiSAAI to predict the interaction probabilities between HER2 and all antibodies targeting protein antigens in the SAbDab.

### 3.5 Interpretability analysis of MultiAAI

When training deep learning models to decipher the intrinsic binding rules between antibodies and antigens, we discovered that, besides the output feature vectors from the Bilinear Attention Network used for classifier training, the attention maps also offer valuable insights. These maps help reveal which positions on the antigen are most likely to interact with the antibody.

The SARS-CoV-2 spike protein in complex with the 4A8 antibody (PDB ID: 7C2L). For visualization, we extracted the antigen chain (Chain A) as well as the heavy (Chain H) and light (Chain L) chains of the antibody (**Fig.5a)**. In the antigen chain (A chain), the epitopes includes residues A-H133, A-K137, A-R223, and A-Y225, which are key sites on the antigen sequence involved in antibody binding (Vita, et al., 2019). Among them, A–H133 indicates that the 133rd residue in chain A is a histidine (HIS). As the heavy chain plays the dominant role in binding within this complex, we visualized the interaction attention between the heavy chain and the antigen using a heat map (**Fig.5b**). The x-axis represents antigen residues, and the y-axis represents residues in the heavy chain. Higher attention values are indicated by more intense colors.

**Figure 5.**
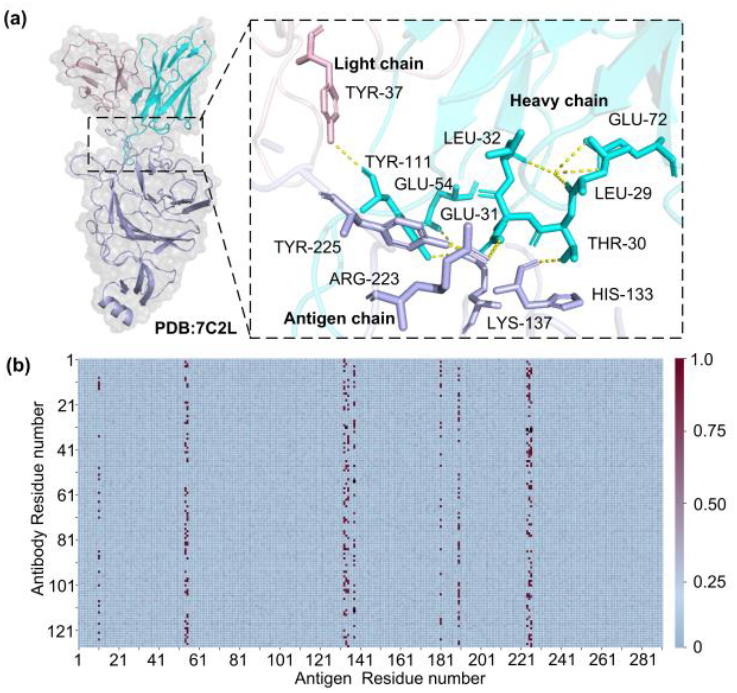
(a) The region of the antibody-antigen complex where interactions occur, with the light chain shown in pink, the heavy chain in cyan, and the antigen chain in purple. (b) The attention heatmap illustrating the antibody-antigen interactions.

Notably, the epitope residues A-H133, A-K137, A-R223, and A-Y225 exhibit strong attention signals, consistent with the biological understanding that only specific residues, rather than the entire sequence, participate in binding. Several key residues in the heavy chain, such as H-T30, H-E31, H-E54, and H-Y111, which interact with the epitope, also show elevated attention scores. Furthermore, the presence of other high-attention regions on the antigen suggests the possibility of alternative epitope recognized by different antibodies. Although this remains a hypothesis requiring further experimental validation, MultiSAAI has indeed provided us with some intriguing insights.

## 4 Discussion and conclusions

Despite the promising performance of MultiSAAI, several aspects of the current work warrant further improvement. Although the Multi-scale Interactive Learner effectively captures the interactions between antibody and antigen sequences, its pursuit of high accuracy comes at the cost of increased computational complexity. Our evaluation indicates that MultiSAAI performs optimally when antigen sequences are shorter than 800 residues; scaling the model to longer antigens while maintaining prediction accuracy remains a challenge. Regarding functional features, although CDR-H3 can be reliably extracted using antibody numbering tools, not all CDR-H3 is directly involved in antigen binding. This potential mismatch introduces noise into the model and may negatively impact classification performance.

In this study, we explicitly considered the distinct contributions of the heavy and light chains, rather than merging or connecting them as usual. In the network architecture, Bilinear Attention Network and multi-fusion convolutional layers were employed to integrate both long-range and local amino acid interactions. In terms of feature selection, residue-level semantic, evolutionary, physicochemical, and geometric features, along with functional features derived from known binding hotspots, provide a comprehensive characterization of antibody–antigen interactions. Importantly, the complementarity among these features ensures low redundancy and high diversity, which contributes to a more informative and robust representation. The experimental results demonstrate that MultiSAAI achieves robust classification performance across both general and target-specific antibody–antigen pairs, suggesting its potential in supporting early-stage drug discovery via large-scale screening.

These findings highlight both the current limitations and strengths of the model. While increasingly complex deep learning architectures may improve prediction accuracy, their effectiveness is often constrained by the limited availability of high-quality labeled data and high computational cost. In this regard, the rapid development of protein language models offers a promising solution by leveraging large-scale unlabeled sequence data to generate informative representations. These pretrained embeddings can enhance feature expressiveness and generalization, particularly under data-scarce conditions. An alternative strategy is to focus on capturing biologically meaningful interaction principles within the deep learning framework. In this context, modeling non-covalent interactions between CDR-H3 and antigenic epitopes presents a promising direction for improving prediction performance in a biologically interpretable and computationally efficient manner.

## 5 Availability

The MultiSAAI server is freely available at http://zhanglab-bioinf.com/MultiSAAI/.

## Supplementary information

### Supplementary Note 1

Dataset Construction for MultiSAAI.

#### (1) Data Extraction

Protein antibody–antigen complexes were collected from the Structural Antibody Database. For each complex, three sequences were extracted: the heavy chain and light chain variable regions of the antibody, and the corresponding antigen sequence. A total of 4440 antibody-antigen sequence pairs were obtained.

#### (2) Redundancy Removal of Antibody Sequences

To prevent highly similar antibody sequences from dominating the dataset, CD-HIT was used to remove redundancy at a sequence identity threshold of 98%. After this filtering step, 2609 unique antibody-antigen pairs remained.

#### (3) Generation of Positive and Negative Samples

To construct a balanced dataset, the 2609 antibody-antigen pairs were first grouped based on antigen similarity. MMseqs2 was used to cluster antigen sequences at 90% identity, forming 747 groups where antigens within a group are highly similar. Positive samples were augmented by random pairing within the group or by swapping antigen partners for antibodies with the same CDR-H3 and belonging to the same group, resulting in an additional 1,181 pairs of positive samples (a total of 3,790 pairs). Negative samples were generated by pairing antibodies and antigens from different groups. To ensure class balance, the number of negative samples per group matched the number of positive samples in that group. This resulted in 3,790 negative pairs. Final dataset comprised 7580 antibody-antigen pairs with a balanced 1:1 ratio of positive to negative samples.

#### (4) Training and Test Set Construction

ClustalW was used to cluster the dataset into nine clusters based on phylogenetic relationships. For five-fold cross-validation, each cluster was split into five equal parts. In each fold, four parts from all clusters were used for training and the remaining part for testing, ensuring that each sample appears in the test set exactly once.

**Figure S1.**
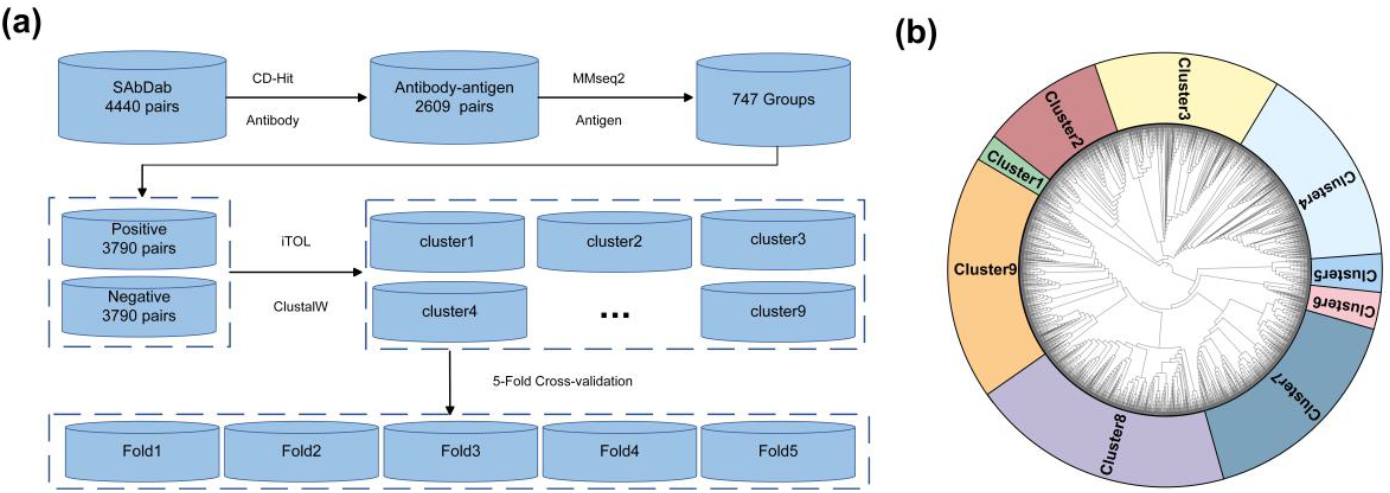
**(a)** The construction process of MultiSAAI’s dataset. **(b)** Generate an evolutionary tree based on the antigen sequence to divide the training set and test set.

### Supplementary Note 2

Detailed description of antibody-antigen functional features.

For the noncovalent interaction between CDR-H3 and epitopes, the functional feature matrix has 5 specific channels, including hydrogen bonding, electrostatic interactions, van der Waals forces, and hydrophobic interactions. Therefore, the shape of the feature matrix (*F*_*eh*_) is *L*_*e*_× *L*_*h*_× 5, Here, *L*_*e*_represents the epitopes length, and *L*_*h*_ represents the CDR-H3 amino acid length.

#### Channel 1

Represents the hydrogen bond interaction potential by taking the minimum between the antibody’s hydrogen bond acceptor value (first value) and the antigen’s hydrogen bond donor value (second value). This channel reflects the limiting factor in hydrogen bond formation.

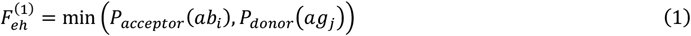

#### Channel 2

Similar to channel 1, but here it takes the minimum between the antibody’s hydrogen bond donor value (second value) and the antigen’s hydrogen bond acceptor value (first value).

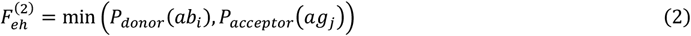

#### Channel 3

This channel quantifies the probability of electrostatic interactions between amino acid pairs. The charge of an amino acid(Q), can be positive, negative, or neutral. To simplify the modeling of electrostatic interactions, only amino acid pairs with opposite charges are considered as potential contributors to the interaction potential.

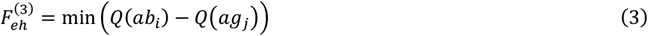

#### Channel 4

Based on the relationship between inter-amino acid distance and interaction strength, we use a Gaussian function to approximate the effect of distance on van der Waals forces. *V*(*x*) denotes the van der Waals volume of an amino acid *x. V*_0_and σ are derived from a Gaussian fit to the aggregate of possible amino acid pair volumes.

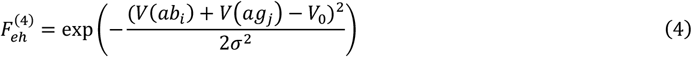

#### Channel 5

We use the hydropathic index to quantify hydrophobic interactions, where *H*(*x*) represents the hydropathic index of amino acid *x. M* denotes the maximum absolute difference in hydropathic index among all possible amino acid pairs

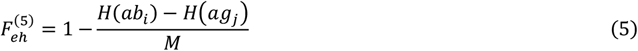

### Supplementary Note 3

Detailed description of the MultiSAAI architecture.

Multi-scale Interactive Learner consists of two components: BAN and MF-CNN, which are used to process the sequence semantic features, physicochemical features, geometric features, and evolutionary features of antibodies and antigens. The CNN block is specifically used to process functional features.

**Figure S2.**
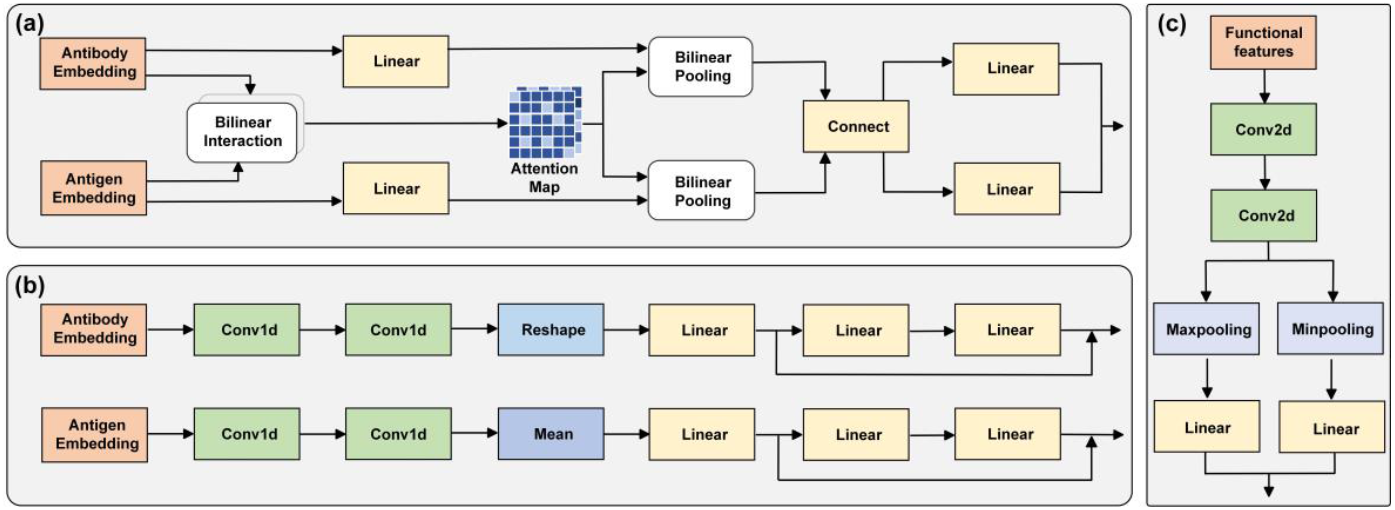
**(a)** Multi-scale Interactive Learner Component 1: BAN. **(b)** Multi-scale Interactive Learner Component 2: MF-CNN. **(c)** CNN block.

### Supplementary Note 4

Detailed parameter settings.

The only difference between MultiSAAI and MultiSAAI_SARS2 in the training process is the epochs. MultiSAAI needs to train 70 epochs, while MultiSAAI_SARS2 only needs to train 50 epochs. It takes 1 day for every fifty epochs.

We use the AdamW optimizer, where the pre-trained model part uses a smaller learning rate (2e-5) to stabilize fine-tuning, while the other network parts use a larger learning rate (1e-4) to speed up training; weight decay (0.0001) is also added to prevent overfitting. The overall strategy aims to fine-tune the pre-trained modules while efficiently training the newly added layers.

**Figure S3.**
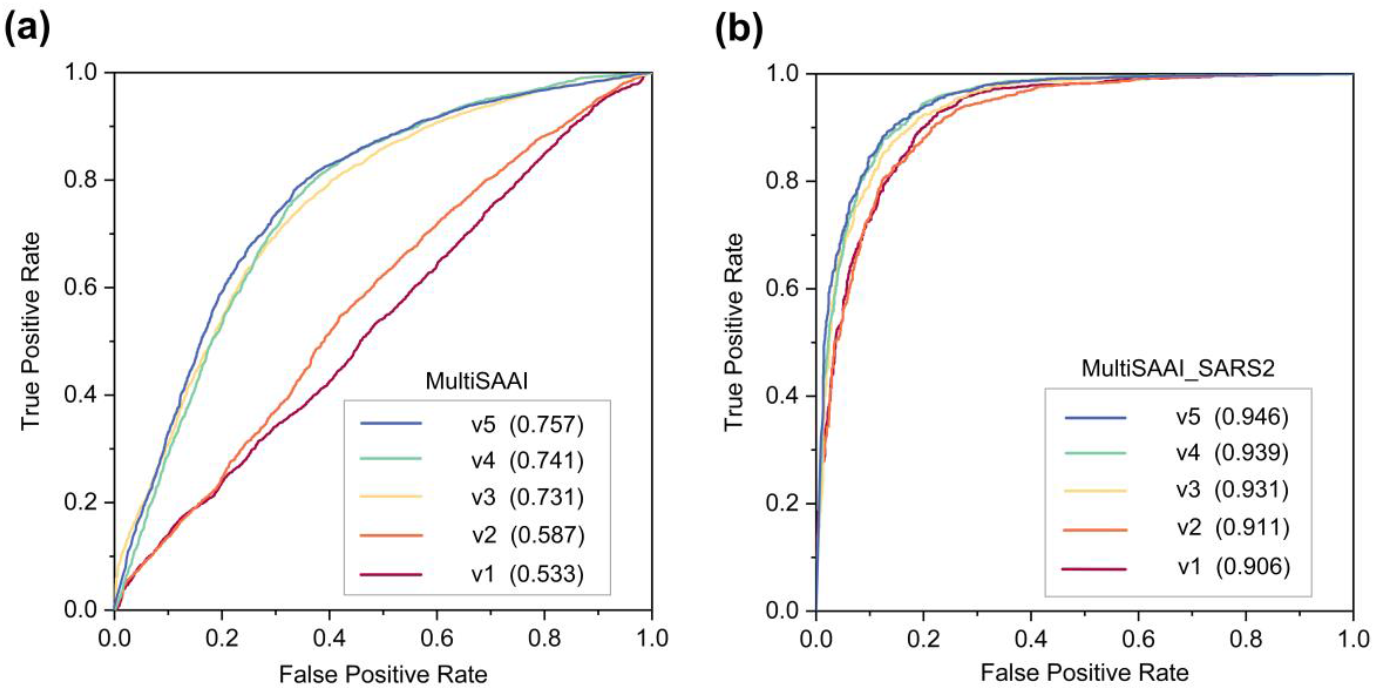
**(a)** Variations in the AUC values across five models for MultiSAAI. **(b)** Variations in the AUC values across five models for MultiSAAI_SARS2.

### Supplementary Note 5

Performance of MultiSAAI and MultiSAAI_SARS2 in ablation study.

## Supplementary Tables

**Table S1.** Detailed description of features.

| Feature | Description | Shape |
| --- | --- | --- |
| **geometric features** | The probability that an amino acid has a helix secondary structure | L×1 |
| **physicochemical features** | A six-dimensional vector representing the steric parameter, polarizability, volume, hydrophobicity, isoelectric point, and sheet probability of each amino acid | L×6 |
| **evolutionary features** | Measures evolutionary similarity between amino acids based on substitution probabilities in aligned sequences. | L×24 |
| **One-hot Encoding** | One-hot encoding represents categorical data by converting each category into a binary vector, where only one position is marked as 1 while all others are 0. In the context of protein sequences, each amino acid is encoded as a unique vector. | L×20 |
| **Normalized Distance to Sequence End** | Indicates a residue's relative position within a sequence, scaled between 0 and 1. | L×1 |

**Table S2.** Performance on the SARS-CoV-2 dataset.

| Method | Precision | Recall | F1 score | AUC |
| --- | --- | --- | --- | --- |
| AbAgIntPre | 0.486 | 0.643 | 0.554 | 0.507 |
| A2binder | 0.825 | 0.852 | 0.837 | 0.924 |
| MultiSAI_SARS2 | 0.862 | 0.885 | 0.873 | 0.946 |

**Table S3.** Performance on the generic dataset.

| Method | Precision | Recall | F1 score | AUC |
| --- | --- | --- | --- | --- |
| AbAgIntPre | 0.502 | 0.709 | 0.587 | 0.507 |
| A2binder_SAbDab | 0.578 | 0.524 | 0.517 | 0.620 |
| MultiSAI | 0.709 | 0.730 | 0.718 | 0.757 |

**Table S4.** Results of five-fold cross validation of A2binder_SAbDab.

| Data | Precision | Recall | F1 score | AUC |
| --- | --- | --- | --- | --- |
| Fold1 | 0.668 | 0.712 | 0.689 | 0.735 |
| Fold2 | 0.545 | 0.805 | 0.649 | 0.543 |
| Fold3 | 0.495 | 0.304 | 0.376 | 0.506 |
| Fold4 | 0.438 | 0.083 | 0.14 | 0.505 |
| Fold5 | 0.745 | 0.714 | 0.729 | 0.809 |
| Average | 0.578 | 0.524 | 0.517 | 0.620 |

**Table S5.**
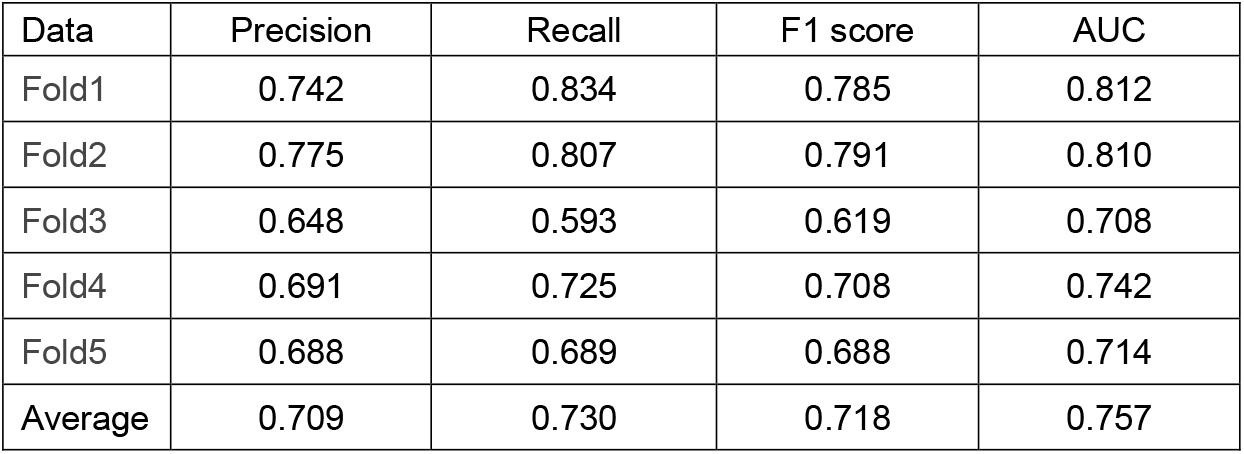
Results of five-fold cross validation of MultiSAAI.

**Table S6.** Ablation experiment results of MultiSAAI_SARS2 (all metrics are the average of five-fold cross validation).

| Model | Precision | Recall | F1 score | AUC |
| --- | --- | --- | --- | --- |
| V1 | 0.815 | 0.834 | 0.824 | 0.906 |
| V2 | 0.807 | 0.866 | 0.836 | 0.911 |
| V3 | 0.830 | 0.873 | 0.851 | 0.931 |
| V4 | 0.846 | 0.877 | 0.861 | 0.939 |
| V5 | 0.862 | 0.885 | 0.873 | 0.946 |

**Table S7.**
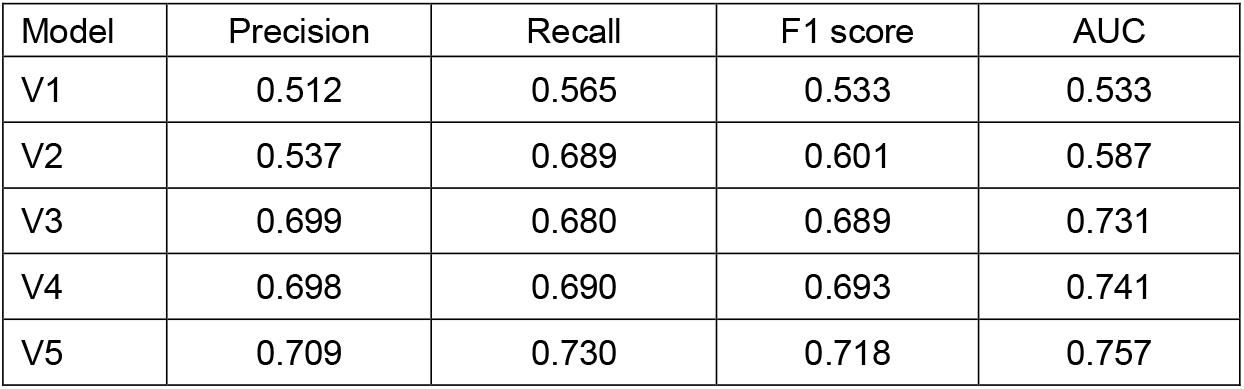
Ablation experiment results of MultiSAAI (all metrics are the average of five-fold cross validation).

| Model | Precision | Recall | F1 score | AUC |
| --- | --- | --- | --- | --- |
| V1 | 0.512 | 0.565 | 0.533 | 0.533 |
| V2 | 0.537 | 0.689 | 0.601 | 0.587 |
| V3 | 0.699 | 0.680 | 0.689 | 0.731 |
| V4 | 0.698 | 0.690 | 0.693 | 0.741 |
| V5 | 0.709 | 0.730 | 0.718 | 0.757 |

**Table S8.**
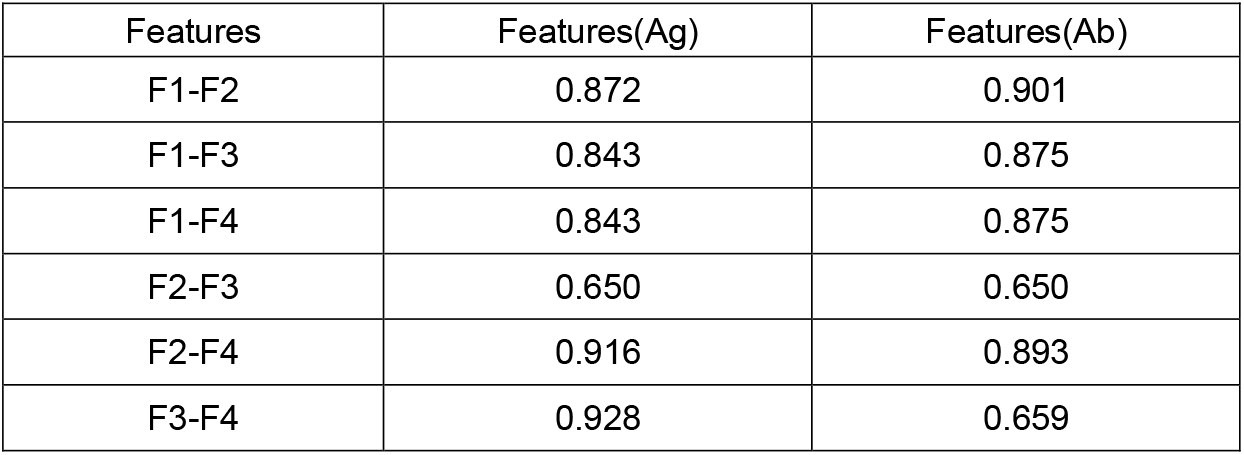
Detailed results on feature complementarity.

| Features | Features(Ag) | Features(Ab) |
| --- | --- | --- |
| F1-F2 | 0.872 | 0.901 |
| F1-F3 | 0.843 | 0.875 |
| F1-F4 | 0.843 | 0.875 |
| F2-F3 | 0.650 | 0.650 |
| F2-F4 | 0.916 | 0.893 |
| F3-F4 | 0.928 | 0.659 |

**Table S9.** Five-fold cross-validation results with and without functional features.

| Data | Without functional features |  |  |  | With functional features |  |  |  |
| --- | --- | --- | --- | --- | --- | --- | --- | --- |
|  | Precision | Recall | F1 | AUC | Precision | Recall | F1 | AUC |
| Fold1 | 0.723 | 0.802 | 0.760 | 0.815 | 0.742 | 0.834 | 0.785 | 0.812 |
| Fold2 | 0.751 | 0.717 | 0.734 | 0.767 | 0.775 | 0.807 | 0.791 | 0.810 |
| Fold3 | 0.64 | 0.697 | 0.615 | 0.672 | 0.648 | 0.593 | 0.619 | 0.708 |
| Fold4 | 0.7 | 0.676 | 0.689 | 0.764 | 0.691 | 0.725 | 0.708 | 0.742 |
| Fold5 | 0.678 | 0.658 | 0.669 | 0.686 | 0.688 | 0.689 | 0.688 | 0.714 |
| Average | 0.698 | 0.69 | 0.693 | 0.741 | 0.709 | 0.730 | 0.718 | 0.757 |

## References

1. Scott, A.M., Wolchok, J.D. and Old, L.J. Antibody therapy of cancer. Nature reviews cancer 2012;12(4):278–287.

2. Andrews, N., et al. Covid-19 vaccine effectiveness against the Omicron (B. 1.1. 529) variant. New England Journal of Medicine 2022;386(16):1532–1546.

3. Watanabe, A., et al. Protective effect of COVID-19 vaccination against long COVID syndrome: A systematic review and metaanalysis. Vaccine 2023;41(11):1783–1790.

4. Steichen, J.M., et al. A generalized HIV vaccine design strategy for priming of broadly neutralizing antibody responses. Science 2019;366(6470):eaax4380.

5. Ledsgaard, L., et al. Basics of antibody phage display technology. Toxins 2018;10(6):236.

6. Grange, R., Thompson, J. and Lambert, D. Radioimmunoassay, enzyme and non-enzyme-based immunoassays. British journal of anaesthesia 2014;112(2):213–216.

7. Kim, J., et al. Computational and artificial intelligence-based methods for antibody development. Trends in pharmacological sciences 2023;44(3):175–189.

8. Pierce, B.G., Hourai, Y. and Weng, Z. Accelerating protein docking in ZDOCK using an advanced 3D convolution library. PloS one 2011;6(9):e24657.

9. Sulea, T., et al. Assessment of solvated interaction energy function for ranking antibody–antigen binding affinities. Journal of chemical information and modeling 2016;56(7):1292–1303.

10. Schneider, C., et al. DLAB: deep learning methods for structure-based virtual screening of antibodies. Bioinformatics 2022;38(2):377–383.

11. Myung, Y., Pires, D.E. and Ascher, D.B. CSM-AB: graph-based antibody–antigen binding affinity prediction and docking scoring function. Bioinformatics 2022;38(4):1141–1143.

12. Gabrielli, E., et al. Antibody complementarity-determining regions (CDRs): a bridge between adaptive and innate immunity. PloS one 2009;4(12):e8187.

13. Adolf-Bryfogle, J., et al. PyIgClassify: a database of antibody CDR structural classifications. Nucleic acids research 2015;43(D1):D432-D438.

14. Vita, R., et al. The immune epitope database (IEDB): 2018 update. Nucleic acids research 2019;47(D1):D339–D343.

15. MacCallum, R.M., Martin, A.C. and Thornton, J.M. Antibody-antigen interactions: contact analysis and binding site topography. Journal of molecular biology 1996;262(5):732–745.

16. Mason, D.M., et al. Optimization of therapeutic antibodies by predicting antigen specificity from antibody sequence via deep learning. Nature biomedical engineering 2021;5(6):600–612.

17. Huang, Y., Zhang, Z. and Zhou, Y. AbAgIntPre: A deep learning method for predicting antibody-antigen interactions based on sequence information. Frontiers in Immunology 2022;13:1053617.

18. Zhang, J., et al. Predicting unseen antibodies’ neutralizability via adaptive graph neural networks. Nature Machine Intelligence 2022;4(11):964–976.

19. Lin, Z., et al. Evolutionary-scale prediction of atomic-level protein structure with a language model. Science 2023;379(6637):1123–1130.

20. Elnaggar, A., et al. Prottrans: Toward understanding the language of life through self-supervised learning. IEEE transactions on pattern analysis and machine intelligence 2021;44(10):7112–7127.

21. Olsen, T.H., Moal, I.H. and Deane, C.M. AbLang: an antibody language model for completing antibody sequences. Bioinformatics Advances 2022;2(1):vbac046.

22. Ruffolo, J.A., Gray, J.J. and Sulam, J. Deciphering antibody affinity maturation with language models and weakly supervised learning. arXiv preprint 2112.07782 2021.

23. He, H., et al. De novo generation of SARS-CoV-2 antibody CDRH3 with a pre-trained generative large language model. Nature Communications 2024;15(1):6867.

24. Olsen, T.H., Boyles, F. and Deane, C.M. Observed Antibody Space: A diverse database of cleaned, annotated, and translated unpaired and paired antibody sequences. Protein Science 2022;31(1):141–146.

25. Deng, J., et al. Nanobody–antigen interaction prediction with ensemble deep learning and prompt-based protein language models. Nature Machine Intelligence 2024;6(12):1594–1604.

26. Zhang, K., Tao, Y. and Wang, F. AntiBinder: utilizing bidirectional attention and hybrid encoding for precise antibody–antigen interaction prediction. Briefings in Bioinformatics 2025;26(1):bbaf008.

27. Kim, J.-H., Jun, J. and Zhang, B.-T. Bilinear attention networks. Advances in neural information processing systems 2018;31.

28. Marks, C. and Deane, C.M. How repertoire data are changing antibody science. Journal of Biological Chemistry 2020;295(29):9823–9837.

29. Dunbar, J., et al. SAbDab: the structural antibody database. Nucleic acids research 2014;42(D1):D1140–D1146.

30. Fu, L., et al. CD-HIT: accelerated for clustering the next-generation sequencing data. Bioinformatics 2012;28(23):3150–3152.

31. Steinegger, M. and Söding, J. MMseqs2 enables sensitive protein sequence searching for the analysis of massive data sets. Nature biotechnology 2017;35(11):1026–1028.

32. Thompson, J.D., Gibson, T.J. and Higgins, D.G. Multiple sequence alignment using ClustalW and ClustalX. Current protocols in bioinformatics 2003(1):2.3. 1-2.3. 22.

33. Liu, Y., et al. Interpretable antibody-antigen interaction prediction by bridging structure to sequence. bioRxiv 2024: 2024.03. 09.584264.

34. Dunbar, J. and Deane, C.M. ANARCI: antigen receptor numbering and receptor classification. Bioinformatics 2016;32(2):298–300.

35. Lefranc, M.-P., et al. IMGT®, the international ImMunoGeneTics information system®. Nucleic acids research 2009;37(suppl_1):D1006–D1012.

36. Meiler, J., et al. Generation and evaluation of dimension-reduced amino acid parameter representations by artificial neural networks. Molecular modeling annual 2001;7(9):360–369.

37. Eddy, S.R. Where did the BLOSUM62 alignment score matrix come from? Nature biotechnology 2004;22(8):1035–1036.

38. Ruby, U. and Yendapalli, V. Binary cross entropy with deep learning technique for image classification. Int. J. Adv. Trends Comput. Sci. Eng 2020;9(10).

39. Rubin, I. and Yarden, Y. The basic biology of HER2. Annals of oncology 2001;12:S3–S8.

40. Scheuer, W., et al. Strongly enhanced antitumor activity of trastuzumab and pertuzumab combination treatment on HER2-positive human xenograft tumor models. Cancer research 2009;69(24):9330–9336.

